# Two novel flavi-like viruses shed light on the plant infecting koshoviruses

**DOI:** 10.1101/2023.02.06.527325

**Authors:** Humberto Debat, Nicolás Bejerman

**Author notes:** Corresponding authors: Humberto Debat,; Nicolás Bejerman.

## Abstract

The family *Flaviviridae* is composed of viruses with a positive sense single-stranded RNA genome and includes viruses that are important veterinary and human pathogens. Most members of the family are arthropod and vertebrate-infecting viruses but more recently flavi-like divergent viruses have been identified in marine invertebrate and vertebrate hosts. The striking discovery of gentian Kobu-sho-associated virus (GKaV) expanded the host range of flaviviruses to plants, which was complemented by a recently reported flavi-like virus in carrot, suggesting they could be grouped in a proposed Koshovirus genus. Here, we report the identification in transcriptomic datasets and characterization of two novel RNA viruses from the flowering plants *Coptis teeta* and *Sonchus asper*, which have genetic and evolutionary affinity to koshoviruses. These two new viruses are members of novel species which were dubbed Coptis flavi-like virus 1 (CopV1) and Sonchus flavi-like virus 1 (SonV1) and with a viral monopartite RNA of ca. 24 kb, likely have the longest genomes among plant-associated RNA viruses yet. Structural and functional annotations of the polyproteins of all koshoviruses resulted in the detection not only of the expected helicase and RNA-dependent RNA polymerase, but also several additional divergent domains such as AlkB oxigenase, Trypsin-like serine protease, methyltransferase, and envelope E1 flavi-like domains. Phylogenetic analysis groups in a monophyletic clade CopV1, SonV1, GKaV and the carrot flavi-like virus robustly supporting the recently proposed genus Koshovirus of plant infecting flavi-like viruses.

---

A massive number of novel viruses are been identified using metagenomic approaches, revealing our limited knowledge about the abundance and diverseness of the global virosphere [1]. This rapid advancement in viromics generates both vast aggregates of new virus data and also challenges in terms of classification [2]. Importantly, it has been established that those newly identified viruses that are detected only from metagenomic data can, should, and have been recognized as novel members when taxonomical proposal are considered by the International Committee on Taxonomy of Viruses (ICTV) [3]. Data mining public sequence datasets has led to the discovery of numerous novel plant viruses. This has resulted in a steady rise in the number of identified viruses, many of which were found in hosts without a previous history of virus infections [4, 5]. Data-driven virus discovery relies on the advancement of open sciences practices, the technical advances of novel sequencing platforms and the huge number of publicly available datasets on the Sequence Read Archive (SRA) of the National Center for Biotechnology Information (NCBI). This resource, which is increasing at an extraordinary rate, includes sequencing data of a large and diverse number of organisms, making the SRA database an invaluable source for virus discovery [6].

The family *Flaviviridae* of small-enveloped viruses is characterized by positive sense single-stranded RNA virus genomes and includes four recognized genera, *Flavivirus, Pestivirus, Pegivirus*, and *Hepacivirus*, and many members are important veterinary and human pathogens. Most of the flaviviruses are arthropod-borne, and are divided into four groups according to their phylogeny and host range [7]. Pestiviruses, pegiviruses and hepaciviruses are vertebrate-infecting viruses that do not require an arthropod vector for transmission [7]. In addition, a vast number of divergent flavi-like viruses have been identified in crustaceans, in free-living parasitic flatworms and decapods as well as the tamanaviruses, that contains viruses from a broad range of vertebrate and invertebrate species [8, 9]. Moreover, strikingly and unexpectedly, two highly divergent flavi-like viruses were identified in plants. The first report came 10 years ago, in a key study unraveling the potential causative agent of a Kobu-sho syndrome in gentian plants, a disease from the mid-80s that led to substantial economic loses in Japan and characterized by stunting, shortened internodes, and tumors on stems, nodes and roots of gentians [10,11]. The surprising affinity of the polyprotein of gentian Kobu-sho-associated virus (GKaV) with flaviviruses and clustering of the virus among - and only - with animal viruses remained baffling for many years. It has been proposed that the origin of these plant virus counterparts could be explained by horizontal virus transfer from invertebrates feeding on plant roots or leaves [12]. Recently, a plant virus infecting carrot was described clustering with GKaV, providing additional support to this clade of viruses which were tentatively grouped as Koshovirus of plant infecting flavi-like viruses [13].

Here, we mined the NCBI-SRA database to search for additional plant-associated flavi-like virus sequences employing the Serratus RNA-dependent RNA polymerase (RdRP) search tool (https://www.serratus.io/explorer/rdrp), which resulted in the identification and assembly of the complete coding sequences of two novel flavi-like viruses linked to flowering plants. The detected viruses were associated to *Coptis teeta* and to *Sonchus asper*, which were tentatively named as Coptis flavi-like virus 1 (CopV1) and Sonchus flavi-like virus 1 (SonV1) (**Table 1**) and likely have the longest genomes and predicted polyproteins among plant-associated viruses.

**Table 1.** Summary of novel flavi-like viruses identified from plant RNA-seq data available in the NCBI database.

| Plant host | Taxa/<br>family | Virus name/<br>Abbreviation | Bioproject ID/<br>Data citation | Length (nt) | Accession<br>number | Protein ID | Length (aa) | Highest scoring virus- protein/E-<br>value/query coverage%/identity% (Blast P) |
| --- | --- | --- | --- | --- | --- | --- | --- | --- |
| Golden thread herb<br>(*Coptis teeta*) | dicot/<br>*Ranunculaceae* | Coptis virus 1/<br>CopV1 | PRJNA433257/<br>[19] | 24205 | BK062902 | Pol | 7716 | CtFLV1-Pol/0.0/82/**36.67** |
| Prickly sow thistle<br>(*Sonchus asper*) | dicot/<br>*Asteraceae* | Sonchus virus 1/<br>SonV1 | PRJNA371565/<br>[20] | 24059 | BK062903 | Pol | 7803 | CtFLV1-Pol/0.0/82/**36.82** |

The pipeline for virus discovery was implemented as described elsewhere [14, 15]. We explored the serratus database employing the serratus explorer tool [16] to identify plant-associated datasets within the SRA which are linked to flavivirus-like sequences. Those SRA libraries that matched the query sequences (alignment identity > 45%; score > 10) were further analyzed in detail. The nucleotide raw sequence reads from each SRA experiment were obtained and processed using the Trimmomatic tool available at http://www.usadellab.org/cms/?page=trimmomatic. The filtered reads were then assembled *de novo* using the rnaSPAdes tool on the Galaxy.org server with standard parameters. The transcripts obtained from the de novo assembly were searched against a collection of flavi-like virus protein sequences available at https://www.ncbi.nlm.nih.gov/protein?term=txid11050[Organism] using bulk local BLASTX searches with an E-value threshold of < 1e-5. Tentative virus-like contigs were then curated through an iterative process that involved mapping the filtered reads using BLAST/nhmmer to extract a subset of reads related to the query contig, extending the contig using the retrieved reads, and repeating the process until the contig was polished. The extended and polished transcripts were then reassembled using the Geneious v8.1.9 alignment tool with high sensitivity parameters. The mean coverage of each assembled virus sequence was calculated using Bowtie2 available at http://bowtie-bio.sourceforge.net/bowtie2/index.shtml with standard parameters. The ORFs of the virus sequences were predicted using ORFfinder (https://www.ncbi.nlm.nih.gov/orffinder/), the presence and architecture of translated gene products were determined using InterPro (https://www.ebi.ac.uk/interpro/search/sequence-search) and the NCBI Conserved Domain Database v3.20 (https://www.ncbi.nlm.nih.gov/Structure/cdd/wrpsb.cgi). In addition, for remote protein homology detection the HHPred tool (https://toolkit.tuebingen.mpg.de/tools/hhpred) was used with the PDB_mmCIF70_10_Jan database. Transmembrane regions were predicted using the TMHMM - 2.0 tool available at https://services.healthtech.dtu.dk/service.php?TMHMM-2.0. The predicted proteins of the assembled viruses were NCBI-BLASTP searched against the non-redundant protein sequences (nr) database and only two virus sequences returned as best hit the polyprotein of the plant infecting carrot flavi-like virus 1 (CtFLV1) and GKaV. All other virus sequences which returned hits to invertebrate flavi-like viruses were discarded. The flavi-like viruses were detected in a library of the flowering plant *Coptis teeta* in a leaf sample obtained in Yunnan, China and in library of a seed sample from France of *Sonchus asper*, and the final assembled virus sequences had a mean coverage of 21.7X (CopV1) and 17.3X (SonV1) (**Table S1**). All additional RNAseq libraries obtained from plants or the genus *Coptis* (84) or *Sonchus* (9) were assessed by read mapping to explore the presence of these viruses in other datasets. While no *Sonchus* libraries were found to have virus reads corresponding to SonV1, four additional libraries of leaf and root samples of both *Coptis teeta* and *Coptis chinensis* from Yunnan and from Nanning China were positive for CopV1, suggesting that this virus could be prevalent (**Table S1**). The assembled consensus sequences corresponding to CopV1and SonV1 were deposited in GenBank under accession numbers BK062902 and BK062903, respectively.

The sequences of CopV1and SonV1 were determined to be 24,205 and 24,059 nt in size, respectively, and they have one single large open reading frame (ORF) coding a 7,716 and 7,803 amino acids (aa) polyprotein, respectively (**Figure 1**). Both assembled viruses presented a highly AU rich short 5’ UTR (CopV1: 143 nt, 74.8% AU; SonV1: 97 nt, 78.4% AU) and a relatively long 3’ UTR (CopV1: 924 nt 52.6% AU; SonV1: 550 nt 55.1% AU). The presence of multiple in-frame stop codons upstream of the proposed AUG start codon of SonV1and the relatively high similarity and synteny of the 5’ terminal region of the SonV1 and CopV1 polyproteins clearly suggest that the assembled viruses are coding-complete. Both encoded polyproteins had the two expected conserved domains reported for GKaV and CtFLV1: a Helicase (HEL, CopV1: pfam00271, 4,100-4,452 aa coordinates, e-value = 2.79e-15; SonV1: pfam00271, 4,207-4,538 aa coordinates, e-value = 2.27e-05) and a RdRP (CopV1: cd01699, 6,940-7,264 aa coordinates, e-value = 3.79e-07; SonV1: cd01699, 7,026-7,349 aa coordinates, e-value = 4.96e-08) (**Figure 1**). In addition, several other domains were detected for the first time in these viruses large polyproteins, some of them also found at equilocal regions in GKaV and CtFLV1, thus probably *bona fide* conserved domains that could suggest structural or functional relevance (**Figure 1**). For instance, a FtsJ-like methyltransferase domain was detected upstream the RdRP region in all four viruses (MeT, CopV1: pfam01728, 6,298-6,522 aa coordinates, e-value = 0.01; SonV1: pfam01728, 6,385-6,609 aa coordinates, e-value = 7.34e-03). This MeT region shared a 75% aa identity among the four viruses. Another domain found in all the viruses was a trypsin-like serine protease immediately upstream of the HEL (Trp, CopV1: IPR009003, 3,898-4,072 aa coordinates; SonV1: IPR009003, 3,989-4,157 aa coordinates). This peptidase region shared a 67% aa identity among the four viruses. Using the HHPred tool a relevant conserved region was found in the four viruses showing significant probability to encode an envelope glycoprotein E of flaviviruses, located at the typically structural region of flaviviruses polyproteins and flanked downstream by transmembrane domains (EnvE1, CopV1: 7LCH_A, 2,173-2,572 aa coordinates, probability = 96.13%; SonV1: 7LCH_A, 2,247-2,652 aa coordinates, probability = 95.66%) (**Figure 1**). It has not escaped our attention that animal enveloped viruses acquire their envelope from the plasma membrane of the host cell and plant cells have cell walls that naturally prevent this exit strategy, thus the detection of flavi-like envelope domains in these plant viruses should be taken cautiously. Perhaps these viruses could replicate via budding from intracellular membranes as other plant enveloped viruses. Further, in SonV1, downstream these transmembrane regions a NS3 protease domain was detected with low confidence but worth mentioning (NS3, 6URV_H Yellow fever protease, 2,958-3097 aa coordinates, probability = 53.6%). Other highly divergent domains found with relatively lower levels of confidence include a nuclease like domain upstream the EnvE1 region, with cues of baculovirus hidrolases (Nuc, CopV1: 7WN7_B, 1,469-1,632 aa coordinates, probability = 95.16%; SonV1: 7WN7_B, 1,556-1,670 aa coordinates, probability = 95.34%). In addition, a Oxoglu/Fe-dep_dioxygenase region was found shared only by CopV1 and SonV1 (AlkB, CopV1: pfam13532, 642-780 aa coordinates, e-value = 3.86e-07; SonV1: pfam13532, 737-873 aa coordinates, e-value = 2.28e-07). The AlkB domain found in some viral genomes appears to be a RNA repair domain that protects the viral RNA genome [17]. Finally, some highly divergent - and unexpected for viruses - domains were detected including a ribonuclease E like region in CopV1 and SonV1 (rE, PRK10811, e-value = 0.03-0.06), a PTZ00121 like region in CtFLV1 and GKaV (PTZ, cl31754, e-value = 0.04-1.83e-03) and an adenylate kinase was found in CopV1 (AK, PRK13808, 966-1,096 aa coordinates, e-value = 0.03). Moreover, six transmembrane domains were identified in the CopV1 polyprotein, and eight in the SonV1 polyprotein, while eight and 12 transmembrane domains were identified in CtFLV1 and GKSaV polyproteins at equilocal-conserved regions, respectively (**Figure 1**). Future studies should complement the predictions described in this report to consolidate the structural and functional landscape of these viruses’ polyproteins. The CopV1 and SonV1 polyprotein shared 63.1% aa identity between them, and both polyproteins shared the highest identity (36.7% and 36.8%, respectively) with that one encoded by CtFLV1. The CopV1 and SonV1 RdRP domain shared 94.6% aa identity, while the helicase (Hel) domain shared 82% aa identity between them. On the other hand, the RdRP domain of both novel viruses shared 83%, 77% and 40-54% aa sequence identity with those of CtFLV1, GKSaV and reported flavi-like arthropod viruses, respectively. In the Hel domain, the corresponding values are 56%, 46% and 25-35%, respectively.

**Figure 1.**
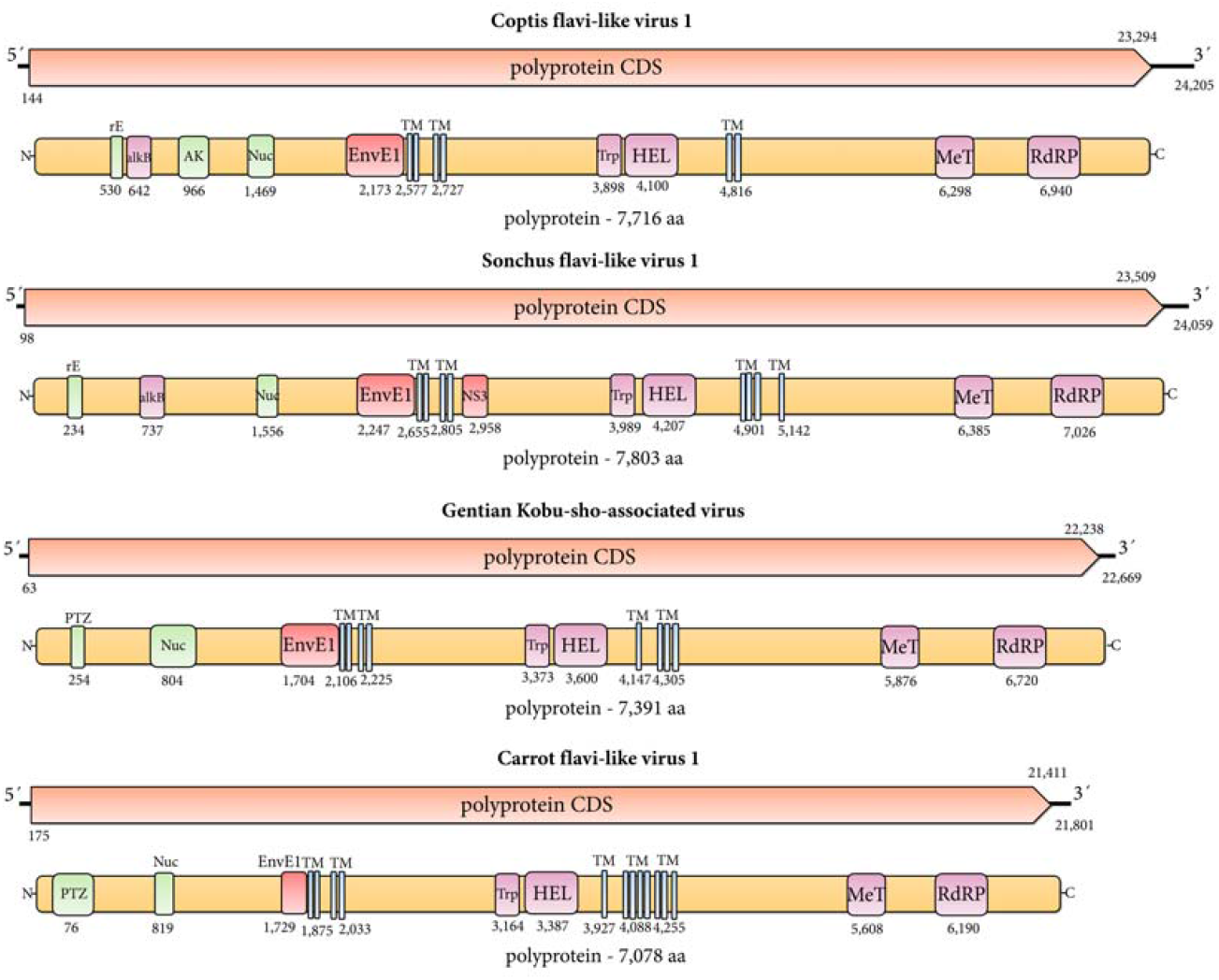
Schematic representation of the genome organization of Coptis flavi-like virus 1 (CopV1), Sonchus flavi-like virus 1 (SonV1), gentian Kobu-sho-associated virus (GKaV) and carrot flavi-like virus 1 (CtFLV1). The predicted coding sequences are shown in orange arrow rectangles. Size in nucleotides, molecular weights in kilo Daltons of predicted proteins and starting coordinates of predicted proteins in aa are indicated. Abbreviations: TM, transmembrane domains; rE, ribonuclease E domain; AK, adenylate kinase; AlkB, Oxoglu/Fe-dep_dioxygenase; Nuc, nuclease domain; EnvE1, envelope glycoprotein E; NS3, NS3 protease; Trp, trypsin-like serine protease; HEL, helicase; MeT, FtsJ-like methyltransferase; RdRP, RNA-dependent RNA-polymerase. Colors of the domains rectangles indicate prediction software: violet, NCBI-CDD; red, HHPred; green, InterPro; light blue, TMHMM.

In order to determine the phylogenetic relationships of CopV1 and SonV1 with other members of the *Flaviviridae* family and flavi-like viruses to unravel its possible evolutionary history, maximum likelihood (ML) phylogenetic analyses were carried out using aa sequences of the RdRp and helicase domains. The RdRp and helicase of the CopV1 and SonV1 viruses were aligned with corresponding sequences of selected members of the *Flaviviridae* family and flavi-like viruses using MAFTT 7 (https://mafft.cbrc.jp/alignment/software/) with the BLOSUM62 scoring matrix and the G-INS-i best-fit algorithm. The aligned sequences were used to construct a maximum likelihood phylogenetic tree in MegaX [18], with 1,000 bootstrap replicates to establish the best-fit model. Phylogenetic analysis showed that CopV1 and SonV1 formed a distinct clade with the plant-associated flavi-like viruses GKSaV and CtFLV1, which was distantly related with the cluster of arthropod flavi-like viruses that have large genomes (**Figure 2A** and **B**). As previously described, both CtFLV1 and GKSaV were found to be associated with plant hosts; CtFLV1 with carrots [13], while GKSaV with gentian and peonies [10, 11]. These flavi-like viruses had the longest genomes among plant-associated viruses to date, were the only flavi-like viruses identified in plants, and have been proposed to be classified as members of a novel genus among the *Flaviviridae* family, provisionally dubbed Koshovirus [13]. CopV1 and SonV1 have a similar genomic organization and are genetically and phylogenetically linked to CtFLV1 and GKSaV, suggesting that CopV1 and SonV1 are two novel members of the Koshovirus genus. Moreover, these two viruses have, by far, the biggest polyproteins among plant-infecting viruses.

**Figure 2.**
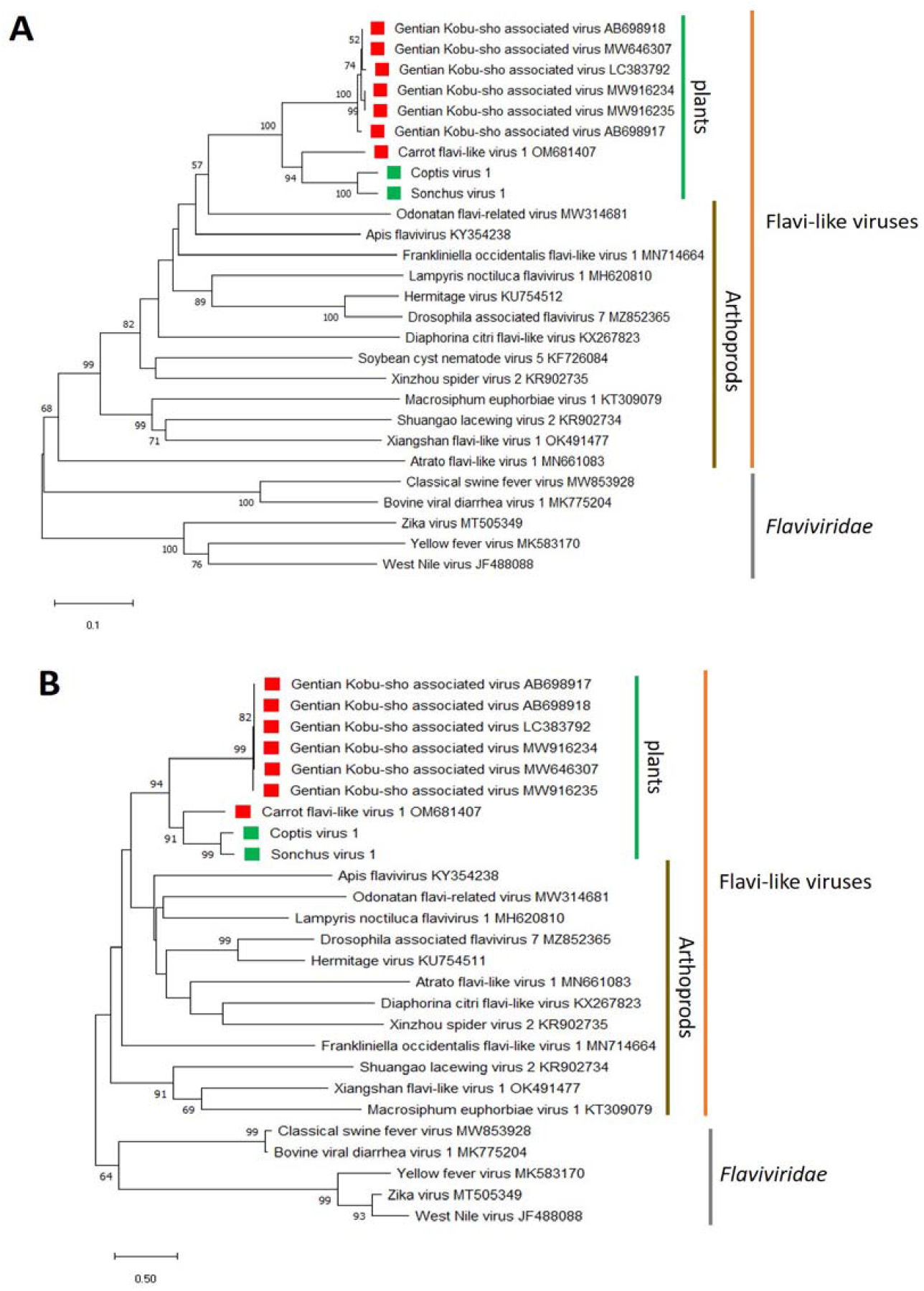
Maximum likelihood phylogenetic trees reconstructed using the conserved RNA dependent RNA polymerase (A) and helicase (B) domains of Coptis flavi-like virus 1 (CopV1), Sonchus flavi-like virus 1 (SonV1) and of representative *Flaviviridae* and flavi-like viruses. Bootstrap values above 50% are shown (1,000 replicates). CopV1 and SonV1 are indicated by green squares, and gentian Kobu-sho-associated virus isolates, as well as carrot flavi-like virus 1 are indicated by red squares. The scale bar the substitution per site.

In conclusion, the analysis of public SRA data provides a valuable resource for identifying novel RNA plant viruses. Our study took a closer look at the plant-infecting Koshovirus and resulted in the discovery of two new plant-associated flavi-like viruses, with the longest genomes among known plant viruses. This discovery doubles the number of known plant-infecting flavi-like viruses, provides clues on additional conserved domains with functional and structural relevance, and strengthens the recent proposal of the genus Koshovirus. Further research is necessary to fully understand the biology and ecology of these viruses, including their prevalence, transmission mechanisms, and potential impact on their hosts.

## Acknowledgments

We would like to express a sincere gratitude to the generators of the underlying data used for this work. By following open access practices and supporting accessible raw sequence data in public repositories available to the research community, they have promoted the generation of new knowledge and ideas.

## Statements and Declarations

### Compliance with ethical standards Ethics approval

This article does not contain any studies with human participants or animals performed by any of the authors.

### Author Contributions

Humberto Debat and Nicolas Bejerman contributed to the study conception and design, data analysis. The manuscript was written by both authors, who commented and reviewed it. All authors read and approved the final manuscript.

### Data Availability

Nucleotide sequence data reported are available in the Third Party Annotation Section of the DDBJ/ENA/GenBank databases under the accession numbers TPA: BK062902-BK062903. The virus sequences are also provided as Supplementary Material 1.

---

**Supplementary Table 1.** Plant RNA-seq datasets assessed for virus mapping available in the NCBI database. Libraries used to generate virus consensus sequences are in bold. Mapping was implemented with Bowtie2 with standard parameters. “Coverage” indicates mean coverage.

| Run | BioSample | Bases | Experiment | geo_loc_name | Organism | Sample Name | Tissue | Virus Reads | Coverage |
| --- | --- | --- | --- | --- | --- | --- | --- | --- | --- |
| SRR6705066 | SAMN08473777 | 7.88 G | SRX3679186 | China:Nanning | *Coptis chinensis* | CcRT2 | root | 0 | 0X |
| SRR6705067 | SAMN08473778 | 6.85 G | SRX3679185 | China:Nanning | *Coptis chinensis* | CcRT3 | root | 413 | 2.1X |
| SRR6705068 | SAMN08473775 | 5.30 G | SRX3679184 | China:Nanning | *Coptis chinensis* | CcLF3 | leaf | 24 | 0.1X |
| SRR6705069 | SAMN08473776 | 7.76 G | SRX3679183 | China:Nanning | *Coptis chinensis* | CcRT1 | root | 0 | 0X |
| SRR6705070 | SAMN08473773 | 8.89 G | SRX3679182 | China:Nanning | *Coptis chinensis* | CcLF1 | leaf | 0 | 0X |
| SRR6705071 | SAMN08473774 | 8.62 G | SRX3679181 | China:Nanning | *Coptis chinensis* | CcLF2 | leaf | 0 | 0X |
| SRR5947132 | SAMN07367940 | 5.40 G | SRX3105522 | China:Yunnan | *Coptis chinensis* | Franch | leaf | 2,117 | 10.9X |
| SRR6705072 | SAMN08473771 | 7.17 G | SRX3679180 | China:Yunnan | *Coptis teeta* | CtRT2 | root | 0 | 0X |
| SRR6705073 | SAMN08473772 | 5.23 G | SRX3679179 | China:Yunnan | *Coptis teeta* | CtRT3 | root | 860 | 4.4X |
| **SRR6705074** | **SAMN08473769** | **5.54 G** | **SRX3679178** | **China:Yunnan** | ***Coptis teeta*** | **CtLF3** | **leaf** | **4,216** | **21.7X** |
| SRR6705075 | SAMN08473770 | 7.18 G | SRX3679177 | China:Yunnan | *Coptis teeta* | CtRT1 | root | 0 | 0X |
| SRR6705076 | SAMN08473767 | 7.96 G | SRX3679176 | China:Yunnan | *Coptis teeta* | CtLF1 | leaf | 0 | 0X |
| SRR6705077 | SAMN08473768 | 7.93 G | SRX3679175 | China:Yunnan | *Coptis teeta* | CtLF2 | leaf | 0 | 0X |
| **SRR5237194** | **SAMN06309740** | **4.50 G** | **SRX2544259** | **France** | ***Sonchus asper*** | **RNA30-06** | **seed** | **4,119** | **17.3X** |
| SRR12816564 | SAMN16427062 | 3.90 G | SRR12816564 | USA: Pennsylvania | *Sonchus asper* | FUS: Ning | un | 0 | 0X |

## References

1. 1. Greninger, AL (2018). A decade of RNA virus metagenomics is (not) enough. Virus Res 244: 218–229. https://doi.org/10.1016/j.virusres.2017.10.014

2. 2. Koonin EV, Dolja VV, Krupovic M, Varsani A, Wolf YI, Yutin N, et al. (2020) Global organization and proposed megataxonomy of the virus world. Microbiol Mol Biol Rev. 84:1–33. https://doi.org/10.1128/mmbr.00061-19

3. 3. Simmonds P, Adams MJ, Benko M, Breitbart M, Brister JR, Carstens EB, et al. (2017) Consensus statement: virus taxonomy in the age of metagenomics. Nat. Rev. Microbiol. 15:161–168. https://doi.org/10.1038/nrmicro.2016.177

4. 4. Bejerman N, Dietzgen RG, Debat H (2021). Illuminating the plant rhabdovirus landscape through metatranscriptomics data. Viruses 13:1304. https://doi.org/10.3390/v13071304

5. 5. Mifsud J, Gallagher R, Holmes E, Geoghegan J (2022). Transcriptome Mining Expands Knowledge of RNA Viruses across the Plant Kingdom. J. Virol. 24:e0026022. https://doi.org/10.1128/jvi.00260-22

6. 6. Lauber C, Seitz S. (2022) Opportunities and Challenges of Data-Driven Virus Discovery. Biomolecules 12:1073. https://doi.org/10.3390/biom12081073

7. 7. Simmonds P, Becher P, Bukh J, Gould EA, Meyers G, Monath T, Muerhoff S, Pletnev A, Rico-Hesse R, Smith DB, Stapleton JT, Ictv Report C (2017) ICTV virus taxonomy profile: Flaviviridae. J Gen Virol 98:2–3. https://doi.org/10.1099/jgv.0.000672

8. 8. Mifsud J, Costa V, Petrone M, Marzinelli E, Holmes E, Harvey E (2023). Transcriptome mining extends the host range of the Flaviviridae to non-bilaterians. Virus Evol. 9:veac124. https://doi.org/10.1093/ve/veac124

9. 9. Parry R, Asgari S (2019). Discovery of novel crustacean and cephalopod flaviviruses: insights into the evolution and circulation of flaviviruses between marine invertebrate and vertebrate hosts. J Virol 93: e00432–19. https://doi.org/10.1128/jvi.00432-19

10. 10. Kobayashi K, Atsumi G, Iwadate Y, Tomita R, Chiba K, Akasaka S, Nishihara M, Takahashi H, Yamaoka N, Nishiguchi M, Sekine KT (2013) Gentian Kobu-sho-associated virus: A tentative, novel double-stranded RNA virus that is relevant to gentian Kobu-sho syndrome. J Gen Plant Pathol 79:56– 63. https://doi.org/10.1007/s10327-012-0423-5

11. 11. Shaffer C, Michener DC, Vlasava NB, Botermans M, Starre J, Tzanetakis IE (2021) First report of Gentian Kobu-sho-associated virus infecting peony in the United States and the Netherlands. Plant Dis. https://doi.org/10.1094/PDIS-06-21-1316-PDN.

12. 12. Dolja VV, Koonin EV (2018). Metagenomics reshapes the concepts of RNA virus evolution by revealing extensive horizontal virus transfer. Virus Res 244:36–52. https://doi.org/10.1016/j.virusres.2017.10.020

13. 13. Schönegger D, Marais A, Faure C, Candresse T (2022). A new flavi-like virus identified in populations of wild carrots. Arch. Virol.167:2407–2409. https://doi.org/10.1007/s00705-022-05544-1

14. 14. Bejerman N, Dietzgen RG, Debat H (2022) Unlocking the hidden genetic diversity of varicosaviruses, the neglected plant rhabdoviruses. Pathogens 11:1127. https://doi.org/10.3390/pathogens11101127

15. 15. Debat H, Garcia ML, Bejerman N (2023). Expanding the repertoire of the plant infecting ophioviruses. bioRxiv. doi: https://doi.org/10.1101/2023.01.27.525910.

16. 16. Edgar RC, Taylor J, Lin V, Altman T, Barbera P, Meleshko D, Lohr D, Novakovsky G, Buchfink B, Al-Shayeb B, et al. (2022). Petabase-scale sequence alignment catalyses viral discovery. Nature 602:142–147. https://doi.org/10.1038/s41586-021-04332-2

17. 17. Bratlie MS, Drabl SF (2005). Bioinformatic mapping of AlkB homology domains in viruses. BMC Genom. 6:1. https://doi.org/10.1186/1471-2164-6-1

18. 18 Kumar S, Stecher G, Li M, Knyaz C, Tamura K (2018). MEGA X: Molecular evolutionary genetics analysis across computing platforms. Mol. Biol. Evol. 35:1547–1549. https://doi.org/10.1093/molbev/msy096

19. 19. He SM, Liang YL, Cong K, Chen G, Zhao X, Zhao QM, Zhang JJ, Wang X, Dong Y, Yang JL, et al. (2018). Identification and characterization of genes involved in benzylisoquinoline alkaloid biosynthesis in Coptis species. Front. Plant Sci. 9:731. https://doi.org/10.3389/fpls.2018.00731

20. 20. Jayasena AS, Fisher MF, Panero JL, Secco D, Bernath-Levin K, Berkowitz O, Taylor NL, Schilling EE, Whelan J, Mylne JS (2017). Stepwise Evolution of a Buried Inhibitor Peptide over 45 My. Mol. Biol. Evol. 34:1505–1516. https://doi.org/10.1093/molbev/msx104

